# Whole-mount smFISH allows combining RNA and protein quantification at cellular and subcellular resolution

**DOI:** 10.1101/2022.10.05.510616

**Authors:** Lihua Zhao, Alejandro Fonseca, Anis Meschichi, Adrien Sicard, Stefanie Rosa

**Author notes:** These authors contributed equally to this work (Lihua Zhao, Alejandro Fonseca).

## Abstract

Multicellular organisms result from complex developmental processes largely orchestrated through the quantitative spatiotemporal regulation of gene expression. Yet, obtaining absolute counts of mRNAs at a 3-dimensional resolution remains challenging, especially in plants, due to high levels of tissue autofluorescence that prevent the detection of diffraction-limited fluorescent spots. *In situ* hybridization methods based on amplification cycles have recently emerged, but they are laborious and often lead to quantification biases. In this article, we present a simple method based on single molecule RNA fluorescence in situ hybridization (smFISH) to visualize and count the number of mRNA molecules in several intact plant tissues. In addition, with the use of fluorescent protein reporters, our method also enables simultaneous detection of mRNA and protein quantity, as well as subcellular distribution, in single cells. With this method, research in plants can now fully explore the benefits of the quantitative analysis of transcription and protein levels at cellular and subcellular resolution in plant tissues.

## INTRODUCTION

Gene expression studies generally require a precise quantification of mRNAs of interest. These studies have commonly used bulk analysis such as RT-qPCR or RNA sequencing approaches. However, these methods do not provide information regarding cellular context and cell-to-cell variability in gene expression. Alternatively, a technique commonly used to study spatial patterns of gene expression is RNA *in situ* hybridization, but this technique is primarily qualitative. Furthermore, none of these techniques provides subcellular resolution. The development of single-molecule RNA FISH (smFISH) has bridged this gap by allowing the detection of individual transcripts with sub-cellular resolution as well as the precise quantification of the number of mRNAs in single cells (Femino *et al.*, 1998; Raj *et al.*, 2008). The use of smFISH has revealed important insights into gene expression, including the presence of large cell-to-cell variability in mRNAs as well as the ability to measure specific gene transcription parameters, such as transcription and degradation rates, burst fractions, and RNA half-life in single cells (Zenklusen *et al.*, 2008; Iyer *et al.*, 2016; Ietswaart *et al.*, 2017; Baudrimont *et al.*, 2017; Duncan and Rosa, 2018).

In plants, smFISH was first applied to root meristem squashes of *Arabidopsis thaliana* (hereafter referred to as Arabidopsis) (Duncan *et al.*, 2016). Plant tissues have very particular optical properties that are often challenging for the imaging process (Donaldson, 2020). Thus, smFISH in plants was initially applied on tissues with low autofluorescence levels, and with the loss of tissue morphology required to obtain monolayers of cells. Therefore, there is currently still an unmet need for quantitative analysis of mRNA expression with high resolution within intact plant tissues. While smFISH allows specific and quantitative analysis of gene transcription, it lacks information about the final gene products – proteins. While such information could in principle, be acquired by combining mRNA detection with protein immunofluorescence, the existing protocols can be difficult to perform because they require sequentially hybridizing and imaging of mRNAs and proteins (Nehmé *et al.*, 2011; Bayer *et al.*, 2015; Eliscovich *et al.*, 2017; Maekiniemi *et al.*, 2020) or are often not quantitative (Yang *et al.*, 2020).

Here, we present a protocol for smFISH in Arabidopsis whole-mount tissues enabling simultaneous detection of mRNA and proteins with cellular resolution in several intact tissues. To take full advantage of this protocol, we developed a computational workflow to quantify mRNA and protein levels at single-cell resolution. For this, we combined our mRNA and protein imaging with a cell wall stain to precisely assign molecular quantities to specific cells. To illustrate the power of our method, we have estimated the cellular specificity in gene expression using well-known protein reporter lines and determined the subcellular distribution of mRNAs known to be located in specific cellular compartments. With our smFISH whole-mount protocol and image analysis pipeline, we can now quantitatively analyze mRNAs and proteins at the cellular and subcellular levels in plants.

## RESULTS AND DISCUSSION

### smFISH for Arabidopsis whole-mount tissues

High levels of autofluorescence have prevented the detection of single RNA molecules in a broad range of plant tissues (Duncan *et al.*, 2016). These difficulties are further complicated by the fact that smFISH is generally imaged with widefield optical microscopes, which are incompatible with the imaging of thick specimens. Assessing the 3D distribution of RNA molecules implies preserving tissue integrity, optimizing the signal-to-noise ratio, and preventing the fluorescence from out-of-focus layers. The easiest way to overcome the latter is to use confocal microscopy, which allows the collection of optical sections of thick specimens. We, thus, tested whether the classical smFISH protocol would allow the detection of mRNA molecules in intact tissues using confocal imaging. To preserve the morphological integrity of the roots, we embedded the samples in a hydrogel according to Gordillo *et al*. (Gordillo *et al.*, 2020) (**Fig. 1A**) and performed smFISH using probes against the exonic regions of the housekeeping *PP2A* (**Table S1**). Fluorescent spots were visible but the signal-to-background ratio was much lower than for squashed tissues and did not allow for confidently identifying mRNA molecule signals throughout the tissue (**Fig. S1A, B**). We, therefore, included additional clearing steps to further minimize autofluorescence and light scattering, including methanol and ClearSee treatments (Kurihara *et al.*, 2015), which significantly improved the signal-to-noise ratio **(Fig. 1A; Fig. S1 B, C)**. We further confirmed that the signals observed correspond to true mRNA molecules by treatment with RNase A (**Fig. S1C, D**). Next, we added a cell membrane staining step using Renaissance 2200 (Musielak *et al.*, 2016) to allow assigning transcripts to different cells and perform intracellular expression comparisons. In whole-mount root tips, *PP2A* mRNA signals could be observed as punctate dots evenly distributed through the cytoplasm (**Fig. 1C**). As expected, we were able to detect *PP2A* mRNAs across all cell types including differentiated cells within the root (**Fig. 1B, C**). Therefore, absolute mRNA counts can in principle be extracted with whole-mount smFISH (hereafter referred to as WM-smFISH) in connection with positional information and cell identities.

**Figure 1.**
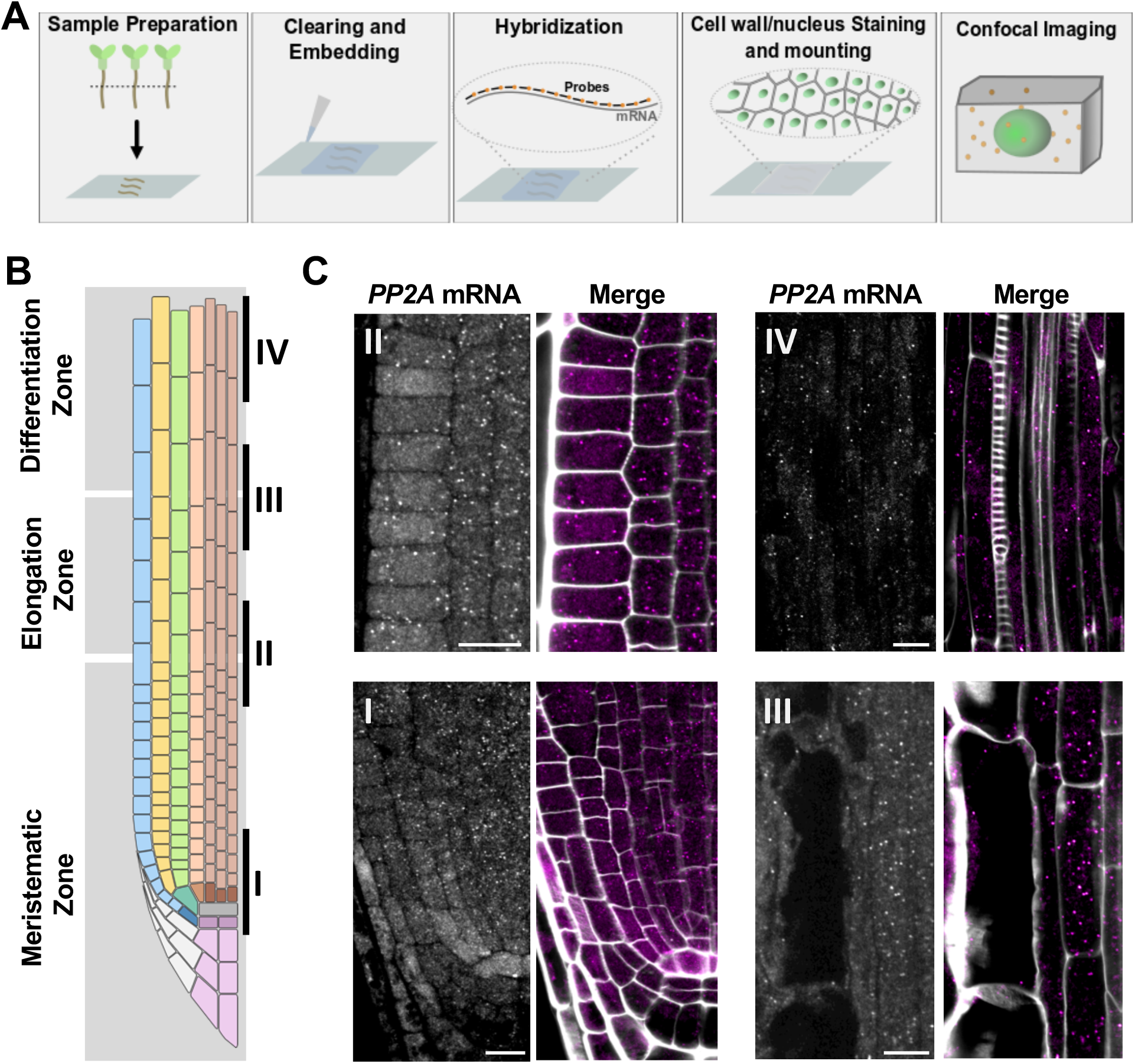
Single-molecule fluorescence *in situ* hybridization in Arabidopsis whole-mounts. (A) Schematic diagram of the whole-mount smFISH method. (B) Schematic of the different developmental regions (I-IV) in the Arabidopsis root. (C) Detection of *PP2A* mRNA molecules in Arabidopsis roots. The contours of cells were visualized through cell wall staining with Renaissance 2200. Scale bars, 10μm.

We then tested whether this method can be applied to various other tissues, including young leaves, inflorescence meristem, ovules, and embryos (**Fig. S2**). We detected *PP2A* mRNA molecules in all tissues analyzed. However, we found a much lower number of mRNAs in these tissues, which is in line with the known expression levels of *PP2A* in different organs (**Fig. S3**). We also observed much weaker signals in leaves and inflorescence (**Fig. S2**). This low signal-to-noise ratio may be caused by the high autofluorescence levels of these tissues. These results demonstrate that single mRNAs can now be detected on several whole-mount Arabidopsis tissues with high specificity and resolution. Spatially quantifying gene expression in highly autofluorescent tissues may nevertheless require further optimizations, such as using different fluorophores, additional clearing steps, or increasing the number of fluorophores per mRNA molecule.

### Simultaneous detection and quantification of mRNA and protein at single cells

While smFISH can provide precise and quantitative measurements of gene expression, it lacks information at the protein level. To that end, we thought to combine WM-smFISH with the detection of fluorescent reporter proteins. We designed probes that targeted the mRNA of VENUS fluorescent protein (**Table S2**), with the aim to simultaneously detect the protein and transcripts expressed by the same transgene (**Fig. 2A**). As a proof-of-concept, we analyzed the auxin signaling reporter line *pDR5rev∷3xVENUS-N7* and a reporter line for the NAC transcription factor *CUP-SHAPED COTYLEDON* 2, *pCUC2∷3xVENUS-N7*, both of which have been extensively characterized (Heisler *et al.*, 2005; Gordon *et al.*, 2007) (**Fig. 2B, E and G**). We choose to perform this analysis on reporter constructs containing three concatenated fluorescent reporters to improve the signal and allow the detection of mRNAs in ‘green tissues’ such as leaves and the inflorescence meristem. Indeed, ninety fluorescent probes are expected to bind these transgenes (*3xVENUS*) as opposed to 48 for the *PP2A* transcripts, which should increase the signal-to-noise ratio by a factor of ~2. We first examined the detection of mRNA and protein in the whole-mount Arabidopsis young leaves, floral primordia, ovule, embryos and roots (**Fig. 2B, E, G; Fig. S4**). The signal-to-noise ratio improved significantly, and mRNA dots could now be easily visualized even in leaves and inflorescence tissues (**Fig. 2B, C, E, G; Fig. S4**). Importantly, the fluorescence of the reporter is well-preserved throughout the WM-smFISH procedure allowing cellular comparison of mRNA and protein distribution (**Fig. 2B-C**).

**Figure 2.**
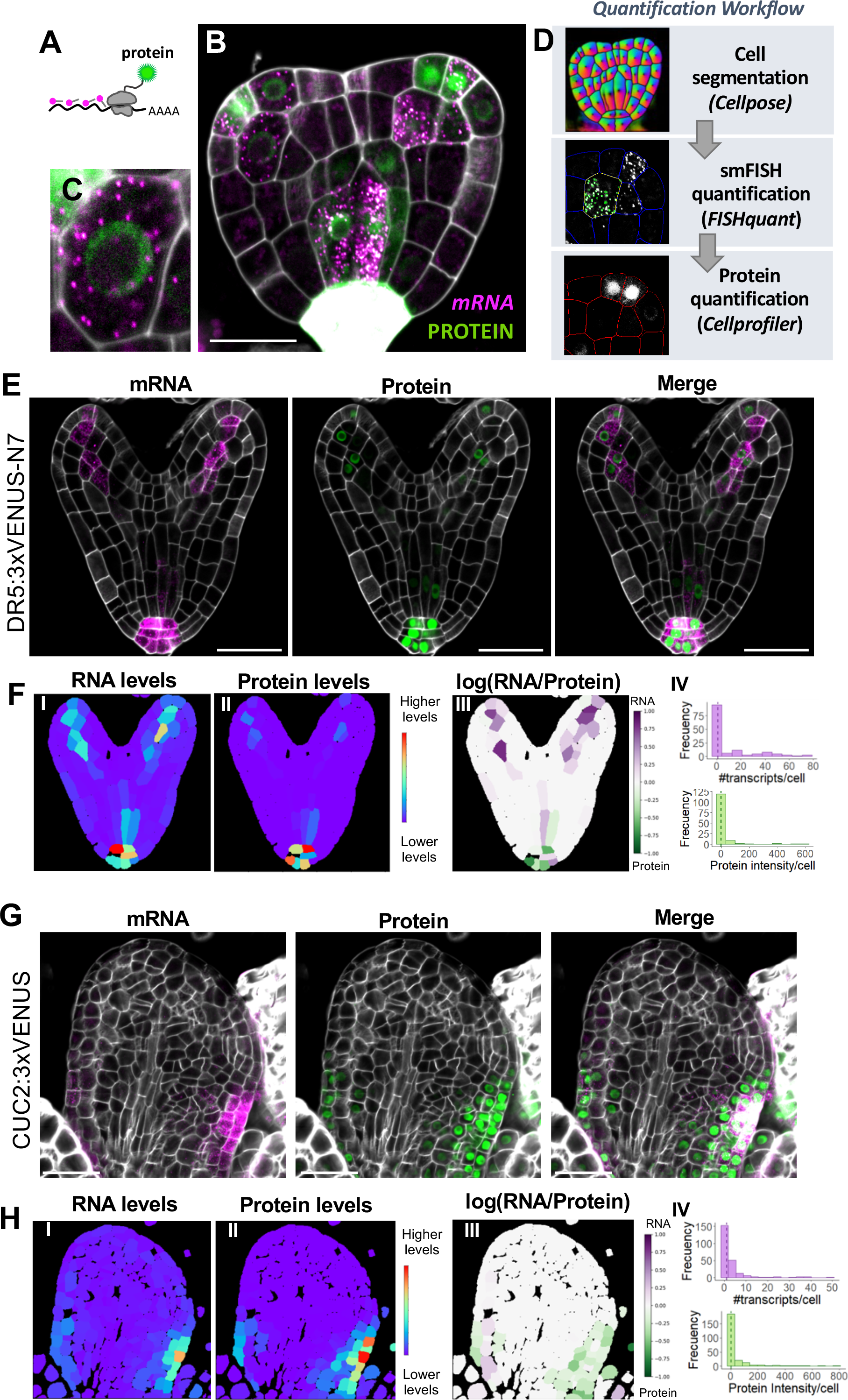
Whole-mount smFISH enables combining RNA and protein quantification. (A) Schematic diagram for simultaneous RNA and protein detection. *VENUS* mRNAs are hybridized and detected with smFISH probes and the VENUS proteins are detected directly through protein fluorescence. (B) Transition embryo expressing *pDR5rev∷3xVENUS-N7* showing detection of VENUS mRNA (magenta) and protein (green). (C) Close-up of a single cell from the embryo presented in B, showing individual mRNAs as single spots and VENUS fluorescence in the nucleus. (D) Workflow diagram showing the three-stepped pipeline for quantitative analysis of wholemount-smFISH with fluorescent protein detection. (E-H) Simultaneous mRNA and protein detection in (E) heart stage embryo and (G) leaf using *pDR5rev∷3xVENUS-N7* and *pCUC2∷3xVENUS-N7* reporter lines, respectively. Confocal microscopy images for mRNA (magenta), protein (green), and merged signals. (F, H) Quantification results for mRNA and protein in heart stage embryo (F)and leaf (H). (I-II) Heatmaps representing the levels of the mean signal intensity per cell detected in each channel (for RNA or protein detection). (III) Heatmap representing the ratio between the RNA and protein signal intensities per cell. (IV) Histograms showing the distribution of the number of transcripts (magenta) or total protein intensity (green) per cell, the median value is indicated with a dashed line. The contours of cells were visualized with Renaissance 2200 dye. Scale bars, 20μm.

To appreciate the spatial differences in the distribution of mRNAs within tissues, we developed a computational workflow to quantify mRNA dots with cellular resolution using WM-smFISH images (**Fig. 2D**). In brief, it segments confocal images based on cell wall (SR2200 dye) signal using Cellpose (Stringer *et al.*, 2021) then uses these cell outlines to estimate the number of mRNA foci per cell using FISH-quant (Mueller *et al.*, 2013; Imbert *et al.*, 2021) and measures the protein intensity fluorescence with CellProfiler (Stirling *et al.*, 2021). A colour scale reflecting intensities (for protein and RNA levels) was finally used to label the segmented cells throughout the confocal images (**Fig. 2F-I, F-II, H-I, H-II**). To visualize the variation in the ratio between mRNA molecules and protein accumulation, we generated heatmaps with the log ratio between the intensity of WM-smFISH and VENUS fluorescent signals (**Fig. 2F-III, H-III**). Here, we choose to use fluorescence intensity rather than the number of mRNA molecules to compare similar measurements. To do this, we first verified that the fluorescence levels per cell correlate with the number of transcripts (**Fig. S5A-B**). The resulting distribution heatmap allows a quantitative and spatial visualization of expression and protein distribution patterns. Histograms can also be used to plot the number of transcripts and protein levels per cell in multiple samples (**Fig 2F-IV, H-IV**).

To validate our quantification workflow, we first measured the number of *PP2A* mRNA molecules in root samples treated with RNAse. As expected, in RNAse treated samples the majority of cells did not show any *PP2A* transcripts, confirming that this pipeline specifically quantified mRNA foci (**Fig. S1, C-E**). Also, our automated detection and counting gave a similar distribution of transcripts per cell as the manual counting of mRNA dots in squashed roots (**Fig. S6G**). Next, we asked whether different image acquisition modes could affect the detection of mRNA dots. We obtained similar distributions for the number of transcripts per cell with widefield and confocal microscopes, as well as with squashed roots and whole-mounts (**Fig. S6H, I**). These results prove that the whole-mount protocol does not compromise the detection and quantification of mRNA molecules. Furthermore, we estimated the cellular specificity of our quantification pipeline by correlating mRNA counts with the level of the corresponding protein in the *pDR5rev∷3xVENUS-N7* line (**Fig. S7**). The VENUS protein levels and mRNA counts were significantly correlated (Pearson R^2^ = 0.3955) (**Fig. S7C**), contrasting with the lack of correlation between the number of *PP2A* transcripts and VENUS protein intensity per cell (Pearson R^2^ = 0.0354) (**Fig. S7D**). Overall, these results validate the accuracy and specificity of our quantification method and indicate that this automated workflow is a useful tool to compare mRNA and protein distributions. Therefore, this approach will be useful to model the transcription/translation dynamics, assess intercellular protein or RNA movement, and analyze co-localization between mRNA and proteins.

We applied our approach to investigate the expression of the *VENUS* reporters at the protein and mRNA levels in different tissues (**Fig. 2F, H; Fig. S7A; Fig. S8**). Globally the spatial distribution of the mRNA molecules and protein signals throughout the tissues were in good agreement with each other and followed the known expression pattern for the two reporter constructs (Heisler *et al.*, 2005; Gordon *et al.*, 2007; Hasson *et al.*, 2011; Hu *et al.*, 2021). We nevertheless did not observe a full expression overlap between the mRNA and protein signals in all tissues. For instance, in the embryo, we observed several cells with high mRNA/protein ratios, this often occurs in cells that express low levels of mRNA (**Fig. 2F; Fig. S5C**). RNA detection by WM-smFISH may therefore be more sensitive than reporter protein imaging. One possible interpretation is that, at this developmental stage, the auxin response has been newly activated in these cells such that the reporter proteins have not yet been translated. Similar discrepancies were also observed in leaf and inflorescence tissues (**Fig 2G-H, Fig S5D, Fig. S8**). For instance, in the young leaf of the *pCUC2∷3xVENUS-N7* line, the reporter proteins appear distributed in more cells than the mRNA molecules (**Fig 2H**). *pCUC2∷3xVENUS-N7* mRNAs appear in cells along leaf margins and may be more consistent with CUC2 function in leaf serration patterns. The diffusion of fluorescent proteins to the neighboring cells seems unlikely due to the high molecular weight of the three concatenated fluorescent proteins and the presence of seven Nuclear Localization Signals. Therefore, these differences are likely to be linked to reporter proteins’ stability considerably exceeding mRNA stability. In this way, the protein signal could persist within a cell even when transcription is not taking place. In dividing tissues such as young leaves and inflorescence meristem, reporter protein distribution could further be extended through cell division, while mRNA molecules would mostly remain in transcriptionally active cells. These results illustrate that fluorescent reporters’ imaging can be combined with WM-smFISH to provide quantitative information on gene activity, with the latter delivering a closer view of the spatial distribution of gene transcription.

### Quantification of mRNA and protein levels with cellular resolution in response to an exogenous stimulus

We further tested our method by analyzing the expression profile of *pDR5rev∷3xVENUS-N7* in Arabidopsis roots in response to the exogenous application of the synthetic auxin naphthalene-1-acetic acid (NAA). In this experiment, we used two different concentrations (1μM and 10μM), to evaluate the difference in sensitivity between WM-smFISH and fluorescent reporter imaging and determine if we could measure quantitative differences in transcript accumulation. A dose-dependent induction in RNA and protein levels was observed (**Fig. 3A-C, E, F**). Globally we observed a coordinate increase in protein and mRNA in the QC and stele cells. However, mRNA signals increased in the epidermis and cortex cells without any apparent activation of the reporter protein fluorescence. The quantification of the mRNA levels or protein fluorescence intensity per cell further confirms a higher increase of mRNA compared to protein at lower NAA concentrations (**Fig. 3B-F**). These results are, therefore, consistent with WM-smFISH being more sensitive. They also demonstrate that combining WM-smFISH with reporter protein imaging can provide quantitative spatio-temporal information on the transcriptional-translational dynamics of gene expression. Combining these measurements with positional information and 3D cell atlas (Montenegro-Johnson *et al.*, 2015; Jackson *et al.*, 2019; Vijayan *et al.*, 2021; Strauss *et al.*, 2022) could provide powerful tools to assess the influence of cellular context on gene expression and translation at a fine scale.

**Figure 3.**
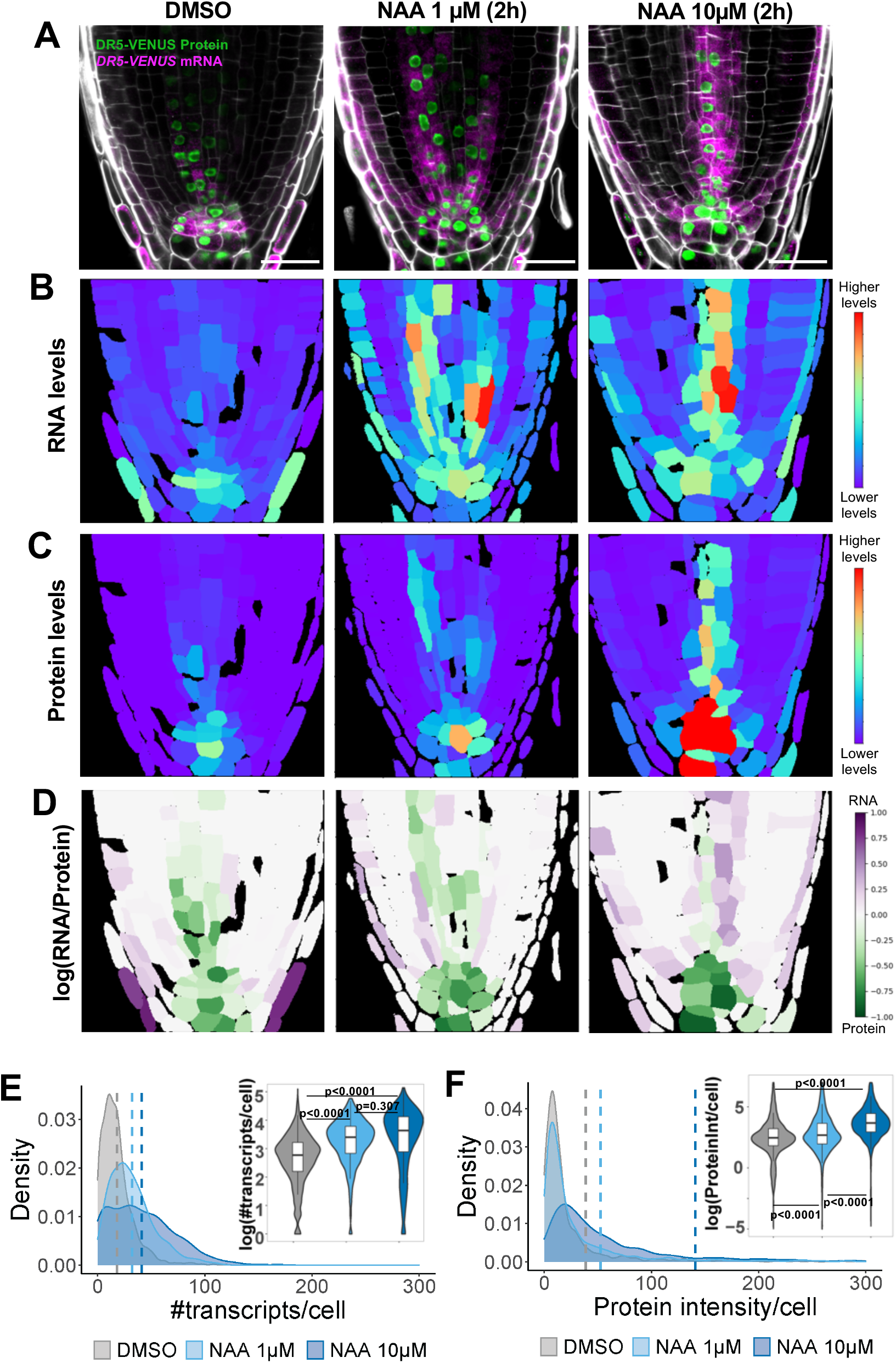
Whole-mount smFISH enables spatial and quantitative characterization of gene expression at RNA and protein levels upon exogenous stimulus. (A) Representative images for the detection of VENUS mRNA (magenta) and protein (green) in *pDR5rev∷3xVENUS-N7* reporter seedlings treated with DMSO, NAA 1 μM, or NAA 10 μM for two hours. The contours of cells were visualized with Renaissance 2200 dye. Scale bars, 20μm. (B-C) Heatmaps representing the levels of the mean signal intensity per cell detected in the channels for (B) RNA or (C) protein detection in the representative images shown in panel A. (D) Heatmaps representing the ratio between the RNA and protein signal intensities per cell in the representative images shown in panel A. (E) Correlation between the number of transcripts per cell area and the mean protein intensity per cell (log-scaled) detected for each representative image shown in panel A. A linear model regression was calculated and the determinant coefficient (R^2^) and adjusted p-value are included in each plot. (F-G) Density plots showing the distributions for (F) the number of transcripts and (G) total protein intensity per cell detected in all the treated roots (DMSO: 1659 cells, NAA 1 μM: 1830 cells, NAA 10 μM: 1456 cells). The dashed lines represent the mean values for each condition. Violin plots showing the log-normalized distributions and p-values for ANOVA/TukeyHSD tests are shown in the inserted panel.

### Subcellular detection and co-localization of mRNA and protein

Subcellular localization of RNAs is important to regulate biological processes, allowing them to find their target, control their translation, or regulate their stability (Martin and Ephrussi, 2009; Das *et al.*, 2021). One example is the mRNAs of nucleoporins (NUP1/NUP2), which are localized and translated next to the nuclear envelope to ensure the proper delivery of the proteins to the nuclear pore complex in yeasts (Lautier *et al.*, 2021). We tested if WM-smFISH can be used to quantitatively evaluate mRNA subcellular localization patterns by colocalizing mRNA spots with fluorescent protein signals. For this, we adapted our automated workflow to segment the protein signal and quantify the number of mRNAs colocalizing with the reporter protein. Using our workflow, we examined the subcellular distribution of *NUP1* mRNA in the apical meristem of Arabidopsis roots expressing NUP1-GFP (**Fig. 4A**). We used probes directed against the *GFP* mRNA to asses the mRNA position which we compared with the localization of the nuclear envelope using NUP1-GFP signal (**Fig. 4A**). As a control, we performed WM-smFISH using *PP2A* probes which we have previously shown to be evenly distributed throughout the cytoplasm (Duncan *et al.*, 2016) (**Fig. 4B**). We detected a significantly higher (*p*<0.0001, Student’s t-test) number of *NUP1* transcripts colocalizing with NUP1-GFP protein compared to *PP2A* (**Fig. 4C**). Nevertheless, we also observed a slightly higher number of *NUP1-GFP* transcripts per cell compared to *PP2A* (*p*<0.0141, Student’s t-test) (**Fig. 4D**). We, therefore, normalized the number of transcripts colocalizing with NUP1-GFP signal by the total number of mRNAs per cell to ensure that indeed a higher proportion of transcripts colocalized with NUP1-GFP. On average, 45.4% of the *NUP1* mRNAs colocalized with NUP1-GFP whereas only 28.7 % of *PP2A* transcripts are present within the nuclear envelope (**Fig. 4E**). The differences (*p*<0.0001, Student’s t-test) indicate that *NUP1* mRNA is preferentially targeted to the nuclear envelope and that WM-smFISH is well suited to investigate the subcellular localization of RNAs and visualize their colocalization with protein partners.

**Figure 4.**
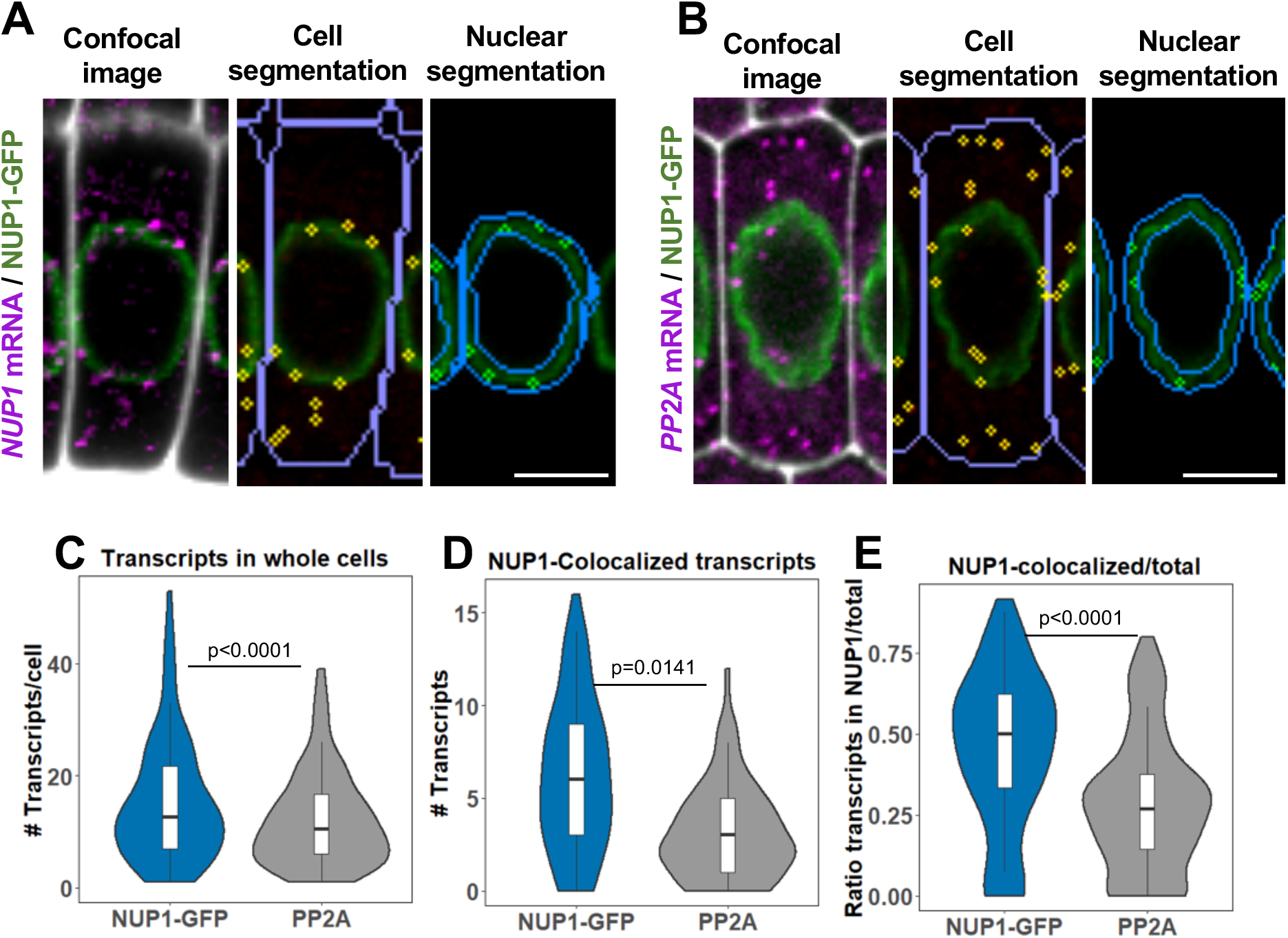
RNA detection by smFISH can be combined with protein detection for subcellular colocalization analysis. (A-B) Representative images to evaluate the subcellular localization of (A) *NUP1-GFP* or (B) *PP2A* mRNAs in cells from the meristematic zone in *NUP1-GFP* expressing roots. Confocal images show the simultaneous detection of the respective mRNA (magenta), NUP1-GFP protein (green), and contours of cells from the Renaissance 2200 dye (white) (left panel). Cells and nuclear envelope were segmented based on Renaissance 2200 and NUP1-GFP signals, respectively. The detected RNA molecules are highlighted in yellow either in the whole cell (middle panel) or colocalizing with the NUP1-GFP signal (right panel). Scale bars, 20μm. (C-E) Violin plots showing the number of *NUP1-GFP* or *PP2A* mRNA molecules per cell (NUP1-GFP: 97 cells, PP2A: 141 cells). A t-test was performed to compare both mRNAs, the p-value is indicated on the graph. The plots show: (C) the number of transcripts per cell, (D) the number of transcripts colocalizing with the NUP1-GFP signal, and (E) the ratio between the number of colocalized transcripts and the total number of transcripts per cell.

In conclusion, we have developed a whole-mount method that enables us to apply smFISH in a variety of intact plant tissues. Determining when and in which tissues and cell types a gene is expressed is essential for their functional characterization. In addition, with the use of fluorescent protein reporters, WM-smFISH can be used for simultaneous detection of mRNA and protein quantity at cellular and subcellular levels. Therefore, this approach will be useful to model the transcription/translation dynamics and for studying regulatory mechanisms associated with developmental and physiological processes. Furthermore, the whole-mount smFISH method presented here may be adapted for use in other plant species and opens up many exciting opportunities for plant researchers.

## MATERIALS AND METHODS

### Plant materials

pDR5rev∷3xVENUS-N7 (N799364) and pCUC2∷3xVENUS-N7 (N23896) were obtained from the Eurasian Arabidopsis Stock Center (uNASC). The NUP1-GFP seeds were a gift from Prof. Chang Liu. All plants were grown as described in the supplementary Materials and Methods.

### Sample preparation

Paraformaldehyde fixed samples were permeabilized and clear through a series of Methanol, Ethanol, and ClearSee (Kurihara *et al.*, 2015) treatment<u>s</u> before being embedded into an acrylamide polymer in which the hybridization was performed. The supplementary Materials and Methods give additional details on the sample preparation and embedding steps.

### *In situ* hybridization

SmFISH probe design and hybridization conditions for different *A. thaliana* tissues are described in Supplementary Materials and Methods.

### Imaging

Whole mount and squashed plant tissues were imaged with a Zeiss LSM800 confocal microscope as described in the Supplementary Materials and Methods.

### Image processing and analysis

Cell segmentations were performed using Cellpose (Stringer *et al.*, 2021). RNA foci were detected and counted using FISH-quant-v3 (Mueller *et al.*, 2013). Co-localisation analysis and heatmap reconstruction were performed using CellProfiler (Stirling *et al.*, 2021). Additional details on the image processing and analyses can be found in the Supplementary Materials and Methods.

## Acknowledgments

We thank Fredric Hedlund and Kathrin Hesse for plant care, and members of the Sicard, and Rosa groups for discussion and comments on the article. We thank Prof. Chang Liu for providing NUP1-GFP seeds.

## Author contributions

LZ, AF, AM, AS, SR designed research. LZ, AM, performed experiments. AF built the image analysis pipeline and performed the quantification analysis. AS and SR wrote the manuscript with help from all authors. All authors agreed on the final version.

## Data availability

Data from this study are not deposited in external repositories but can be requested from the corresponding author.

---

**Figure.S1.**
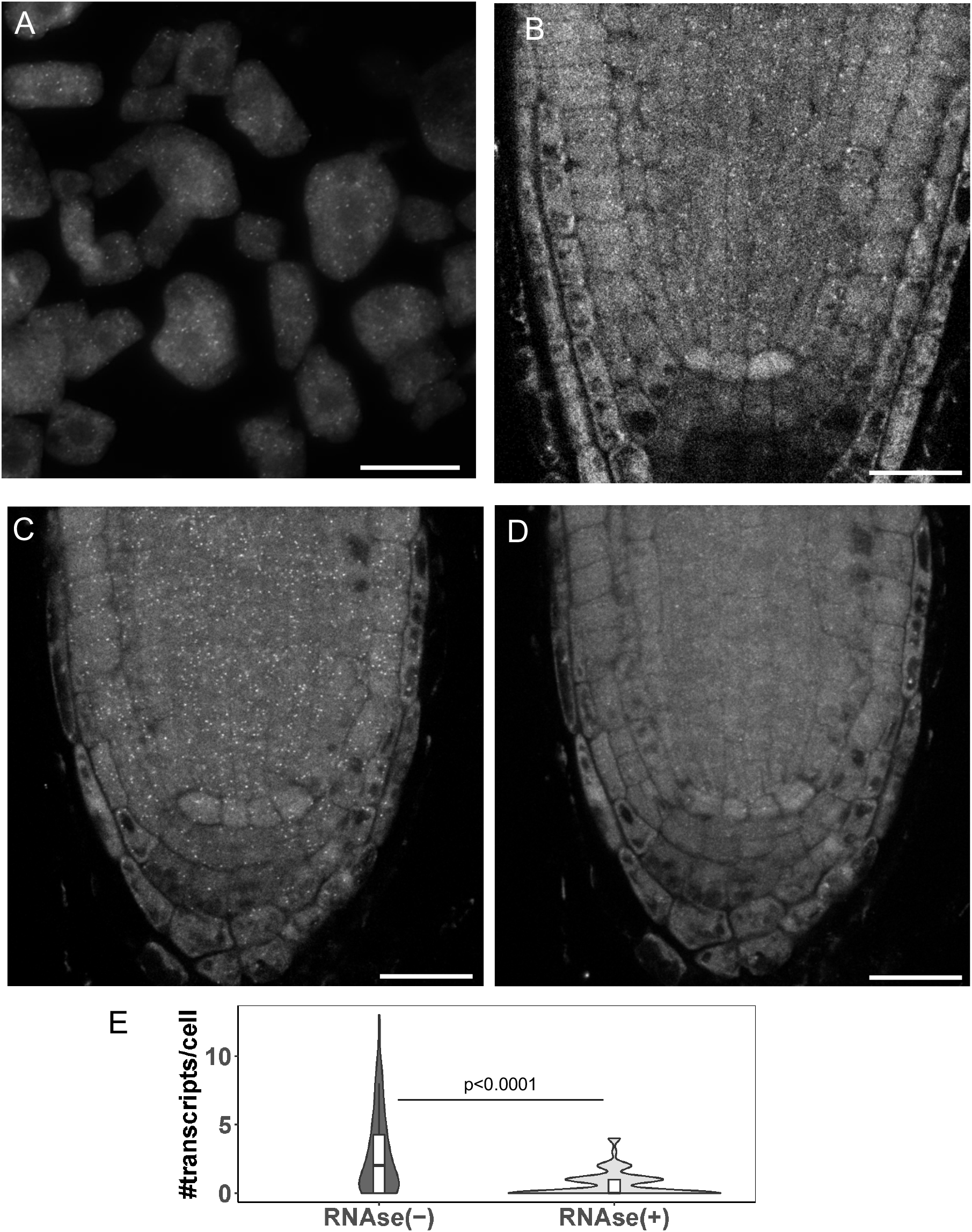
Optimization of clearing procedure for whole-mount smFISH. (A-B) smFISH using probes against *PP2A* mRNA (grey spots). (A) Representative image of smFISH in root meristem squashes of 7-days-old Arabidopsis seedlings. Imaging was performed using a widefield microscope. (B) Whole-mount smFISH in the root meristem of 7-days-old Arabidopsis seedling without ClearSee treatment. Imaging was performed using a confocal microscope. (C) Representative image of whole-mount smFISH method in the root meristem of 7-days-old Arabidopsis seedling after ClearSee treatment. Imaging was performed using a confocal microscope. (D) Whole-mount smFISH of the root depicted in C after 15 minutes of RNase treatment. Scale bars, 20μm. (E) Violin plot comparing the number of transcripts detected before [RNAse(−), panel C] and after RNAse treatment [RNAse(+), panel D]. Boxes inside show the interquartile range (IQR 25-75%), indicating the median values as a horizonal line. Whiskers show the ±1.58xIQR value. A t-test was performed to compare both conditions, the p-value is indicated on the graph. N = 130 cells per condition.

**Fig. S2.**
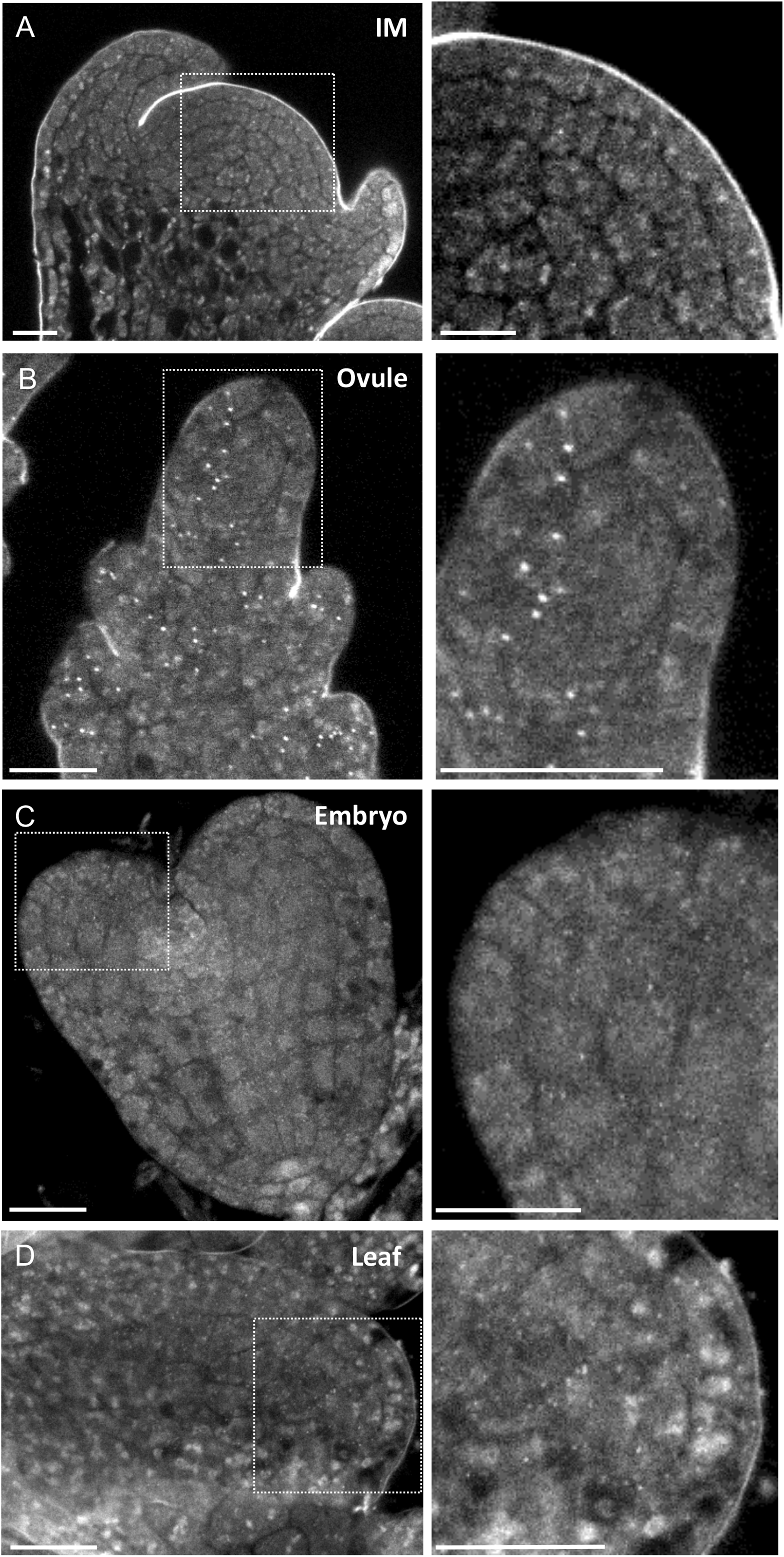
WM-smFISH in other plant tissues using *PP2A* probes. Representative images of whole-mount smFISH in: (A) inflorescence meristem; (B) ovule; (C) embryo; (D) leaf. Right: Zoomed-in images from the regions highlighted with a square on the left panel images. Scale bars, 10μm.

**Fig.S3.**
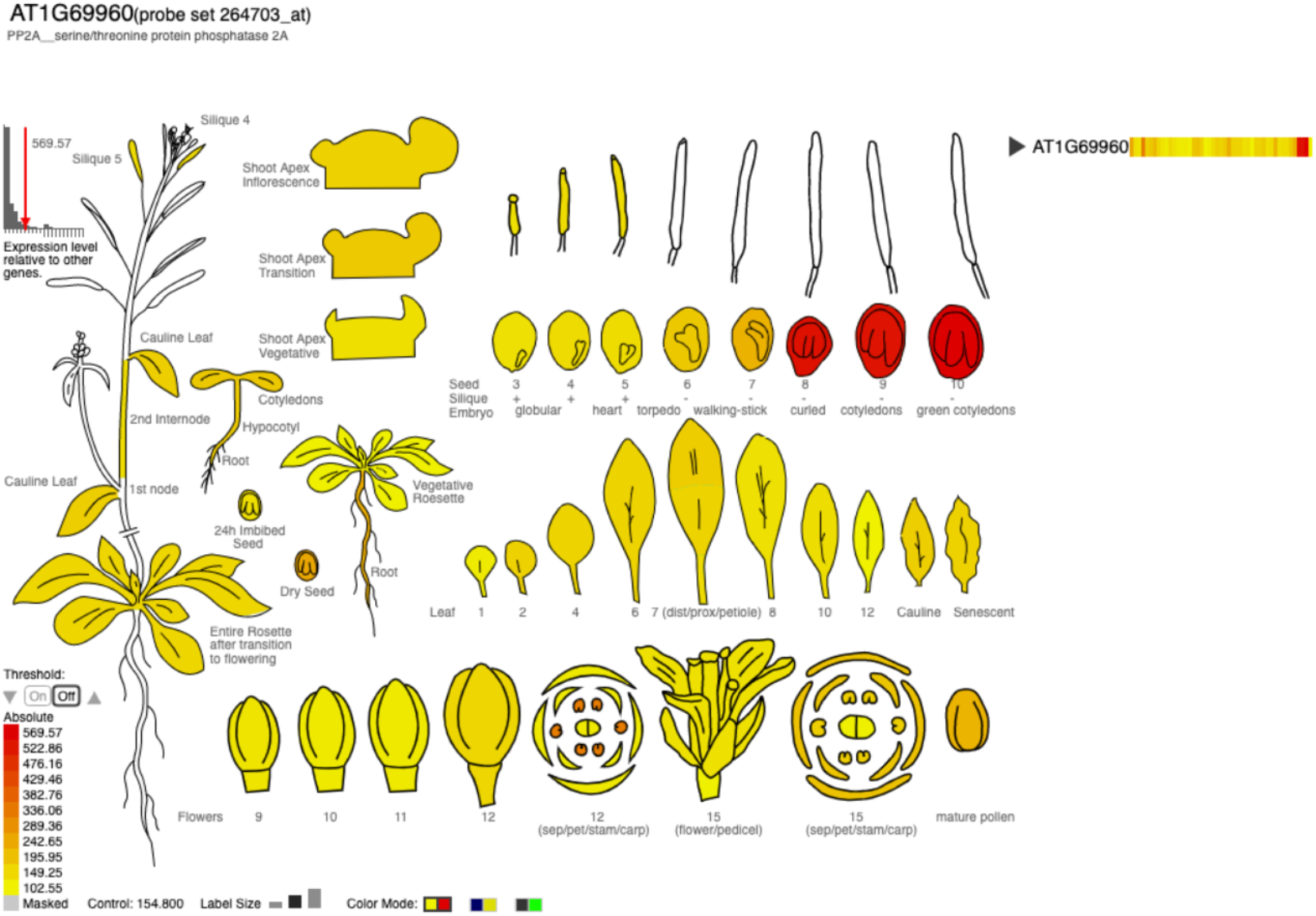
Expression level of *PP2A* in different organs. The above image for gene AT1G69960 (represented by ATH1 probe set 264703_at) was generated using the eFP Browser 2.0 at bar.utoronto.ca by Waese et al 2013.

**Figure.S4.**
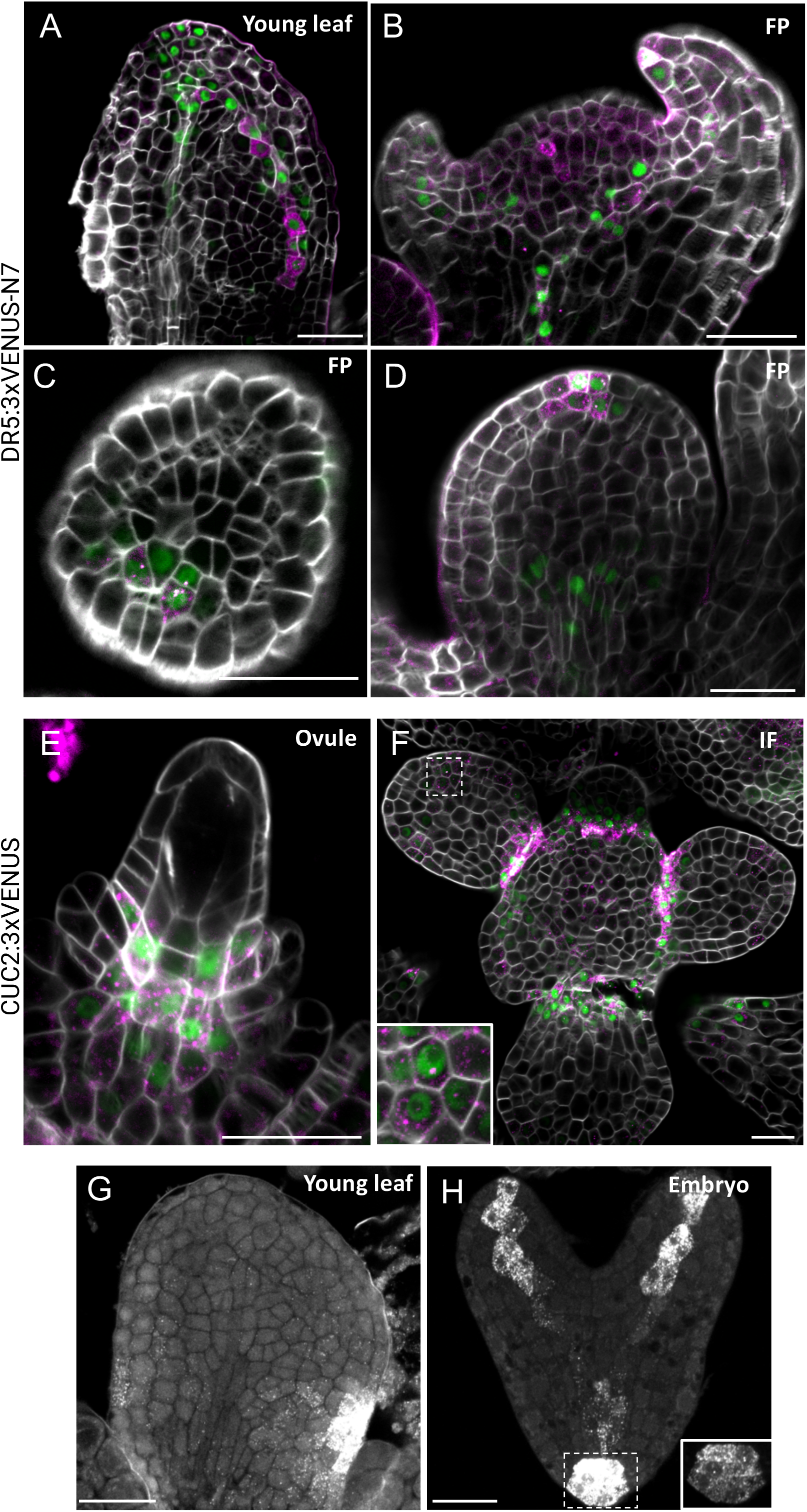
WM-smFISH for simultaneous detection of mRNA and protein in different tissues. (A-F) Representative images for WM-smFISH for the detection of VENUS mRNA (magenta) and protein (green) in *pDR5rev∷3xVENUS-N7* and *pCUC2∷3xVENUS-N7* reporter lines in: young leaf (A); floral primordia (B-D); ovule (E); inflorescence meristem (F). (G-H) WM-smFISH images depicted in Fig. 2E (embryo) and 2G (young leaf) showing only the smFISH probe channel in grey. Inset (H): showing the same region with lower brightness and contrast levels, allowing to observed single dotted signals corresponding to mRNAs. Scale bars, 20μm.

**Fig.S5.**
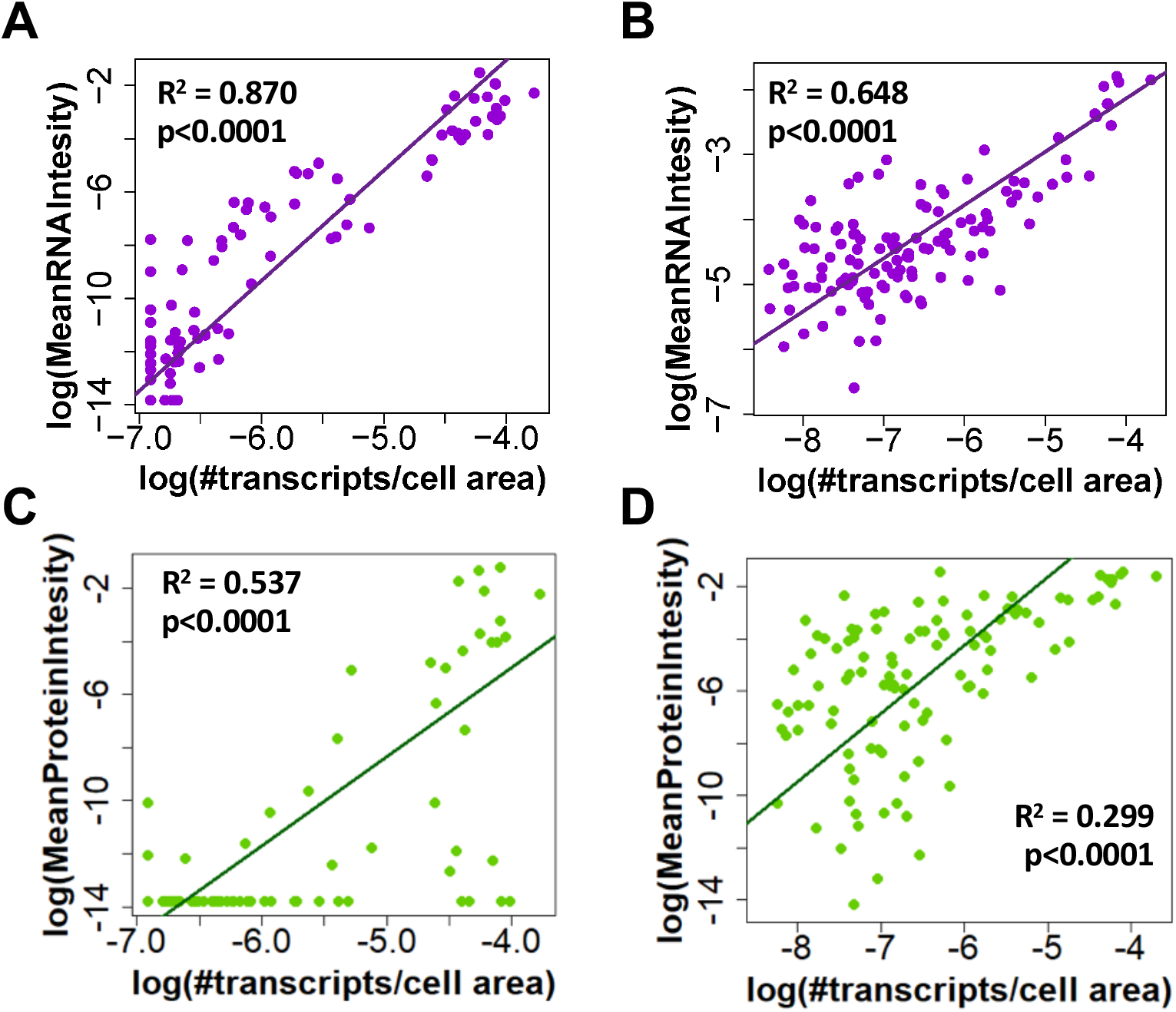
Correlation between the number of transcripts per cell and the total fluorescence intensity of RNA or proteins at the cellular level. Scatter plots showing the correlation between the number of transcripts per cell area and the mean RNA intensity (by smFISH) (A, B) or protein intensity (C, D) per cell (in logarithmic scales) detected for the representative images shown in Figure 2. A linear model regression was calculated and the determinant coefficient (R^2^) and adjusted p-value are included in each plot. (A, C) Heart-stage embryo. (B, D) leaf.

**Figure S6:**
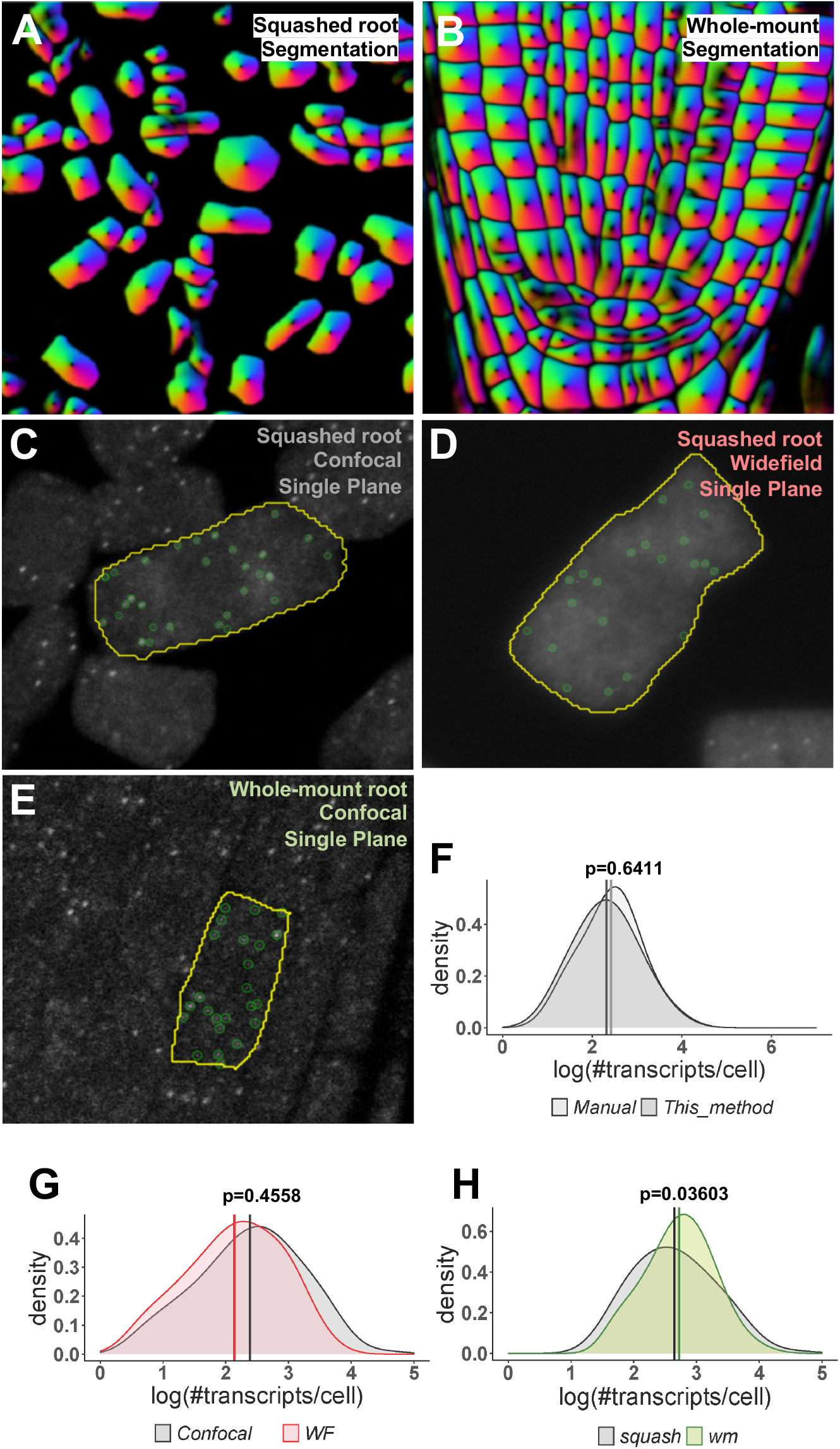
Evaluation of the RNA-single molecule quantification workflow in images acquired by different methods. (A-B) Representative images showing the cell segmentation results obtained for squashed-root tip cells (A) and from a whole-mount root (B) using Cellpose. (C-E) Representative images showing the detected RNA molecules by FISHquant using different acquisition methods: (C) squashed root, confocal microscopy, single plane; (D) squashed root, widefield microscopy, single plane; (E) whole-mount root, confocal microscopy, single plane. (F-H) Density plots comparing the distributions for the number of RNAs per cell using different methods. Distributions were statistically compared using a two-sample centered Kolmogorov-Smirnov test, p-values are indicated on the graph. (F) Comparison between detection by eye (manual) and the workflow from this paper (This_method) in images from a squashed root, obtained with widefield microscopy in Z-planes (N = 61 cells). (G) Comparison between images from squashed roots using confocal or widefield (WF) microscopy (Confocal: 520 cells, WF: 255 cells). (H) Comparison of the distributions obtained from squashed roots (squash) or whole-mount roots (wm) analyzing one plane from confocal microscopy (squash: 520 cells, wm: 1249 cells).

**Fig.S7.**
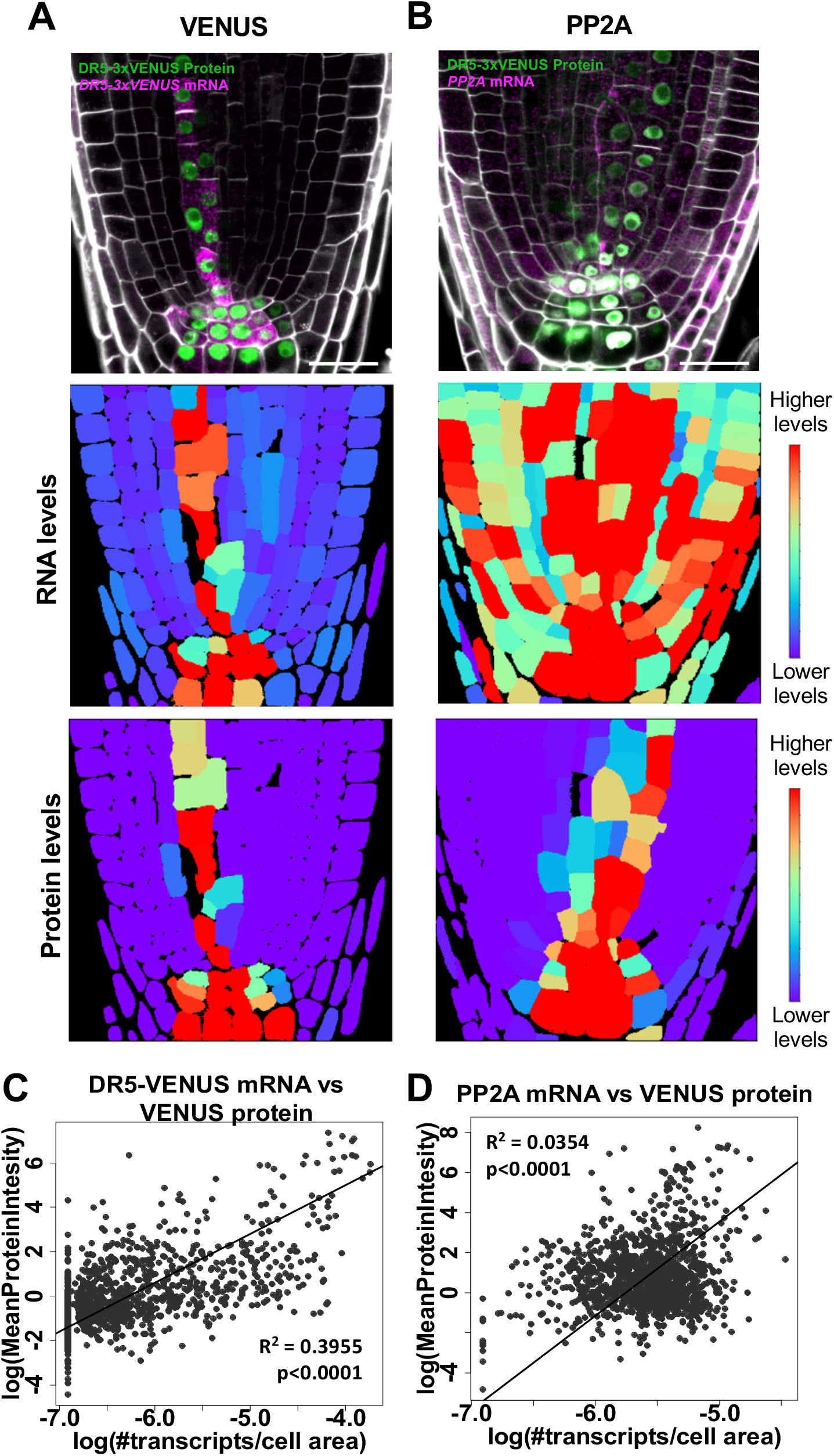
Specificity detection validation for the RNA single-molecule quantification method. Representative images to quantify the VENUS (A) or *PP2A* (B) mRNAs in cells from the meristematic zone in roots from 7-day old *pDR5rev∷3xVENUS-N7* reporter lines. Confocal images (upper panels) show the simultaneous detection of the respective mRNA (magenta), VENUS protein (green), and cell contours with Renaissance 2200 dye (white). Scale bars, 20μm. Heatmaps represent the levels of the mean signal intensity per cell detected in the channels for RNA (middle panels) or protein detection (bottom panels) in the representative images shown in the top panels. Heatmaps represent the ratio between the RNA and protein signal intensities per cell in the representative images shown in the top panels. (C) Correlation between the number of transcripts per cell area and the mean protein intensity per cell (in logarithmic scales) detected for *VENUS* or *PP2A* mRNAs. Linear model regressions were calculated and the determinant coefficient (R^2^) and adjusted p-value are included in each plot (VENUS: 1136 cells, PP2A: 1299 cells).

**Fig.S8.**
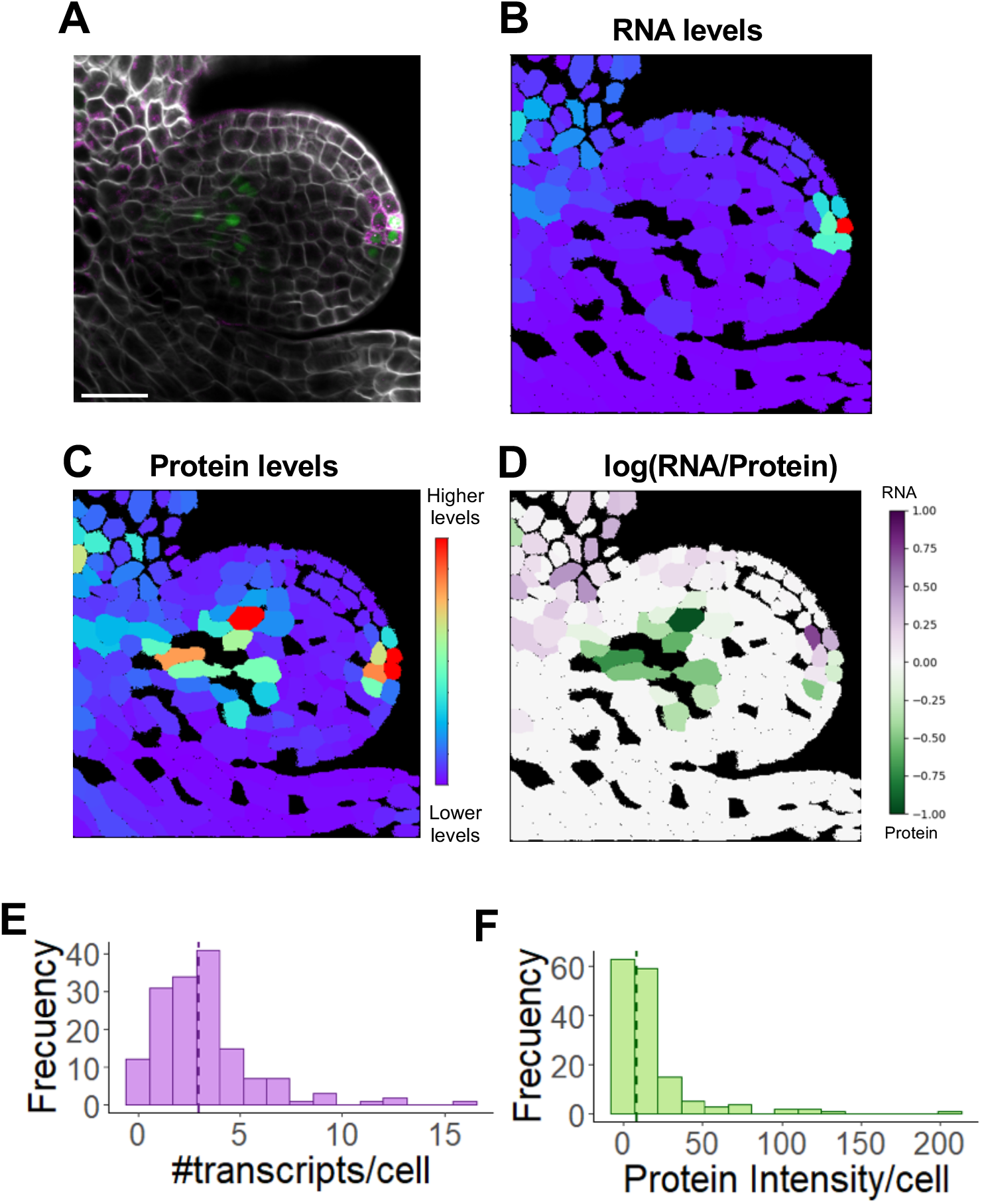
Simultaneous quantification for RNA and protein quantification in floral primordia. Simultaneous mRNA and protein in floral primordia using a *pDR5rev∷3xVENUS-N7* reporter line. (A) Confocal microscopy image detecting mRNA (magenta), protein (green), and cell contours with Renaissance 2200 dye (white). Scale bars, 20μm. (B-C) Heatmaps representing the levels of the mean signal intensity per cell detected in each channel for RNA (B) or protein (C) detection. (D) Heatmap representing the ratio between the RNA and protein signal intensities per cell. (E-F) Histograms showing the distribution of the number of transcripts (E) or total protein intensity (F) per cell (right), the median value is indicated with a dashed line.

